# Bovine serum albumin interaction modulates the cellular potency of the BUB1 inhibitor BAY-1816032: fluorescence, molecular docking, and mesothelioma cell evidence

**DOI:** 10.64898/2026.10.03.756242

**Authors:** Fazel Gorjipour, Ece Çakıroğlu, Şerif Şentürk

## Abstract

Albumin binding can substantially alter the unbound concentration and apparent cellular potency of small-molecule anticancer agents. BAY-1816032 is a selective BUB1 kinase inhibitor with activity in malignant pleural mesothelioma, but its interaction with serum albumin has not been defined. This study combined intrinsic bovine serum albumin (BSA) fluorescence measurements, molecular docking, and cell-viability assays to examine whether albumin association influences BAY-1816032 activity. BAY-1816032 produced pronounced quenching of BSA fluorescence at low drug-to-protein molar ratios, and the quenching response was greater at mildly acidic pH than at mildly basic pH. Docking to BSA (PDB 4F5S) placed the top-ranked pose in Sudlow site I (subdomain IIA), with a Vina score of −7.8 kcal/mol and predicted hydrogen-bond contacts with Tyr149, Arg256, and Asp450. In NCI-H28, NCI-H2052, NCI-H2452, and AB12 mesothelioma cells, supplementation of 10% fetal bovine serum medium with 2% (w/v; 20 mg/mL) BSA shifted BAY-1816032 concentration–response curves to the right. Relative to 10% serum alone, IC□□ values increased from 2.4 to 11.1 µM in H28, 3.6 to 22.0 µM in H2052, 2.0 to 4.9 µM in H2452, and 2.4 to 11.1 µM in AB12. These data support an albumin-dependent reduction in the freely available drug fraction and identify site I as the leading computational binding-site hypothesis. Because fluorescence-intensity and docking data do not independently establish a dissociation constant or quenching mechanism, direct unbound-fraction measurements and competitive displacement experiments are required for quantitative confirmation.

## 1. Introduction

BUB1 is a serine/threonine kinase that contributes to spindle-assembly checkpoint signaling and accurate chromosome segregation. BAY-1816032 is a potent, orally bioavailable BUB1 inhibitor that sensitizes tumor models to taxanes and inhibitors of ATR and PARP [1]. A genome-wide CRISPR screen subsequently identified BUB1 as a druggable vulnerability in malignant pleural mesothelioma (MPM), and pharmacological inhibition with BAY-1816032 suppressed proliferation and anchorage-independent growth in MPM models [2]. These observations motivate development of the compound while also making its extracellular disposition an important determinant of experimental and translational activity.

Serum albumin is the most abundant plasma protein and a major carrier of endogenous ligands and xenobiotics. Reversible albumin association can increase solubility and residence time while lowering the immediately unbound concentration that is available for membrane permeation, target engagement, metabolism, or elimination [3–5]. Albumin contains two principal drug-binding cavities: Sudlow site I in subdomain IIA, classically associated with warfarin, and Sudlow site II in subdomain IIIA, classically associated with ibuprofen [6]. Binding can also vary with pH because albumin conformation and ligand ionization are environmentally sensitive [7].

The rationale for the present work arose from our preceding analytical study of BAY-1816032 in serum. During development of UV, direct-fluorescence, and RP-HPLC–fluorescence assays, quantitative chromatographic recovery required 20% DMSO to release BAY-1816032 from serum proteins, indicating substantial protein association [8]. Because albumin constitutes the major soluble protein fraction of serum, we investigated BSA as a tractable model of this interaction. The analytical characterization and assay validation reported previously are not repeated here; instead, the present study focuses on protein interaction and its functional consequence for cellular potency.

We therefore asked three linked questions: whether BAY-1816032 alters intrinsic BSA fluorescence, whether docking supports a defined albumin-binding pocket, and whether increasing extracellular albumin changes BAY-1816032 potency in mesothelioma cells. The resulting data connect a biophysical interaction with a reproducible rightward shift in cellular concentration–response curves, while preserving a clear distinction between experimental observations and computational predictions.

## 2. Materials and methods

### 2.1 Reagents and study design

BAY-1816032 (MedChemExpress, USA) was dissolved in dimethyl sulfoxide (DMSO) to prepare a concentrated stock solution. BSA and phosphate-buffered saline (PBS) were used for fluorescence experiments. Final DMSO content in protein-containing fluorescence samples did not exceed 1% (v/v). The study comprised (i) intrinsic BSA fluorescence-quenching experiments, including a pH comparison; (ii) structure-based docking to the crystallographic BSA model; and (iii) MTT concentration–response assays under media containing 5% fetal bovine serum (FBS), 10% FBS, or 10% FBS supplemented with 2% (w/v) BSA.

### 2.2 BSA fluorescence measurements

BSA was prepared at 2 µM in PBS. BAY-1816032 was added to final concentrations of 1, 2, 4, 6, 8, and 10 µM, corresponding to BAY-1816032:BSA molar ratios of 0.5:1 to 5:1. Intrinsic protein fluorescence was measured using a Varioskan platform (Thermo Fisher Scientific) with excitation at 280 nm and collection of emission spectra over 300–480 nm. To examine environmental sensitivity, BSA alone and BSA–BAY-1816032 mixtures were additionally compared at pH 7.1 and 7.7. Fluorescence-intensity changes were interpreted qualitatively because absorbance-based inner-filter correction, temperature dependence, fluorescence lifetime, and replicate-level Stern–Volmer analyses were not available for the present dataset.

### 2.3 Protein and ligand preparation for docking

The crystal structure of BSA (PDB ID 4F5S; 2.47 Å resolution) was obtained from the Protein Data Bank [9]. The co-crystallized small molecule, water molecules, and non-essential ions were removed in UCSF Chimera 1.17.1 [10]. Hydrogen atoms were added, non-polar hydrogens were merged, and Gasteiger partial charges were assigned. The protein was minimized using the AMBER force field for 1,000 steepest-descent steps and exported in PDBQT format.

The three-dimensional BAY-1816032 structure was obtained in SDF format from PubChem. Geometry was optimized in Chem3D using MM2 minimization followed by molecular-dynamics refinement. Gasteiger charges and hydrogen treatment were completed in Chimera, rotatable bonds were assigned, and the ligand was exported in PDBQT format.

### 2.4 Molecular docking and interaction analysis

Docking was performed with AutoDock Vina 1.1.2 [11] through PyRx 0.9.2 [12]. The search space enclosed Sudlow site I in subdomain IIA, including the region surrounding the single intrinsic tryptophan residue of BSA (Trp213 in the 4F5S numbering used in the interaction diagram). Default exhaustiveness was used. Poses were ranked by the Vina score, which is reported as a computational scoring estimate rather than an experimentally measured binding free energy or equilibrium constant. Three-dimensional poses were inspected in Chimera and PyMOL; two-dimensional contacts were mapped using LigPlot+ 2.2 [13].

### 2.5 Cell culture and MTT viability assay

Human MPM cell lines NCI-H28, NCI-H2052, and NCI-H2452 and the murine mesothelioma cell line AB12 were maintained in RPMI-1640-based culture medium under standard humidified conditions (37 °C, 5% CO□). For the albumin-modulation experiment, cells were exposed to BAY-1816032 for 72 h in medium containing 5% FBS, 10% FBS, or 10% FBS plus 2% (w/v) additional BSA. The supplemented condition therefore contained 20 mg/mL added BSA, approximately one-half of the typical physiological plasma albumin concentration, in addition to the protein contributed by 10% FBS. Cell viability was determined by MTT reduction using six replicate wells for every BAY-1816032 concentration and medium condition. Separate cell-free blanks were included for each serum/BSA condition, and untreated cells maintained in the corresponding medium served as condition-specific controls. Blank-corrected absorbance values were normalized to the respective untreated controls. Concentration–response curves were fitted by four-parameter logistic (4PL) regression, and IC□□ values were derived from the fitted curves.

### 2.6 Data handling

Concentration–response data were fitted using the four-parameter logistic equation Y = Bottom + (Top − Bottom)/[1 + 10^((LogIC□□ − X) × HillSlope)], where X is the log□□ BAY-1816032 concentration and Y is normalized viability. IC□□ fold changes were calculated as the IC□□ in 10% FBS plus 2% BSA divided by the IC□□ in 10% FBS alone. The six replicate wells at each concentration were blank-corrected and normalized to condition-matched untreated controls before curve fitting. The reported IC□□ values describe the fitted concentration–response shifts; independent biological repetition and confidence-interval-based comparison of complete curves remain desirable for confirmatory inference.

## 3. Results

### 3.1 BAY-1816032 markedly quenches intrinsic BSA fluorescence

BSA displayed the expected broad intrinsic fluorescence emission centered near 330–340 nm after excitation at 280 nm. Addition of BAY-1816032 produced a pronounced decrease in BSA fluorescence, with substantial quenching already evident at the lowest tested drug concentration (1 µM; BAY-1816032:BSA molar ratio 0.5:1). Increasing the drug concentration maintained near-complete suppression of the protein fluorescence signal. The strong response at a substoichiometric molar ratio supports close association between BAY-1816032 and the fluorescent BSA environment, but it does not by itself establish a 1:1 stoichiometry, a binding constant, or a purely static mechanism.

At matched protein and drug conditions, the remaining fluorescence intensity was lower at pH 7.1 than at pH 7.7 (Figure 1). This pattern indicates that the observed interaction and/or the photophysical response is sensitive to modest pH changes. Because pH can alter albumin conformation, ligand protonation, fluorescence quantum yield, and inner-filter effects, the experiment is interpreted as evidence of pH sensitivity rather than a direct measurement of stronger equilibrium binding at acidic pH.

**Figure 1.**
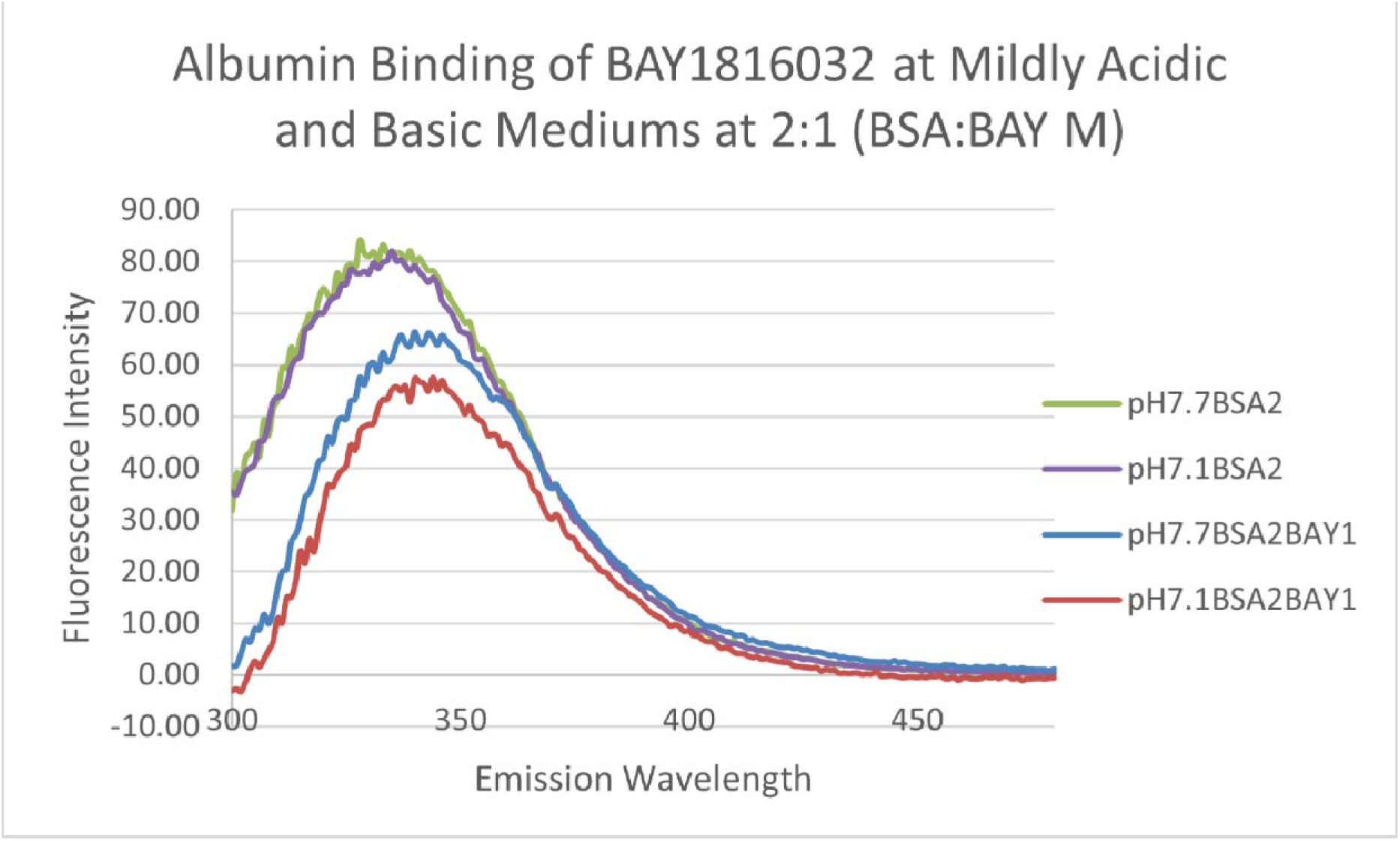
Intrinsic BSA fluorescence in mildly acidic and mildly basic media, with and without BAY-1816032. BSA was used at 2 µM and BAY-1816032 at 1 µM (2:1 BSA:drug molar ratio). The lower residual signal in the pH 7.1 BSA–drug trace indicates pH-sensitive quenching. Excitation was at 280 nm; emission wavelength is shown in nanometers.

### 3.2 Docking places BAY-1816032 in Sudlow site I

The top-ranked docking pose localized BAY-1816032 to Sudlow site I within subdomain IIA and yielded a Vina score of −7.8 kcal/mol. The LigPlot+ map predicted hydrogen-bond contacts with Tyr149 (3.09 Å), Arg256 (2.99 Å), and Asp450 (3.10 Å), together with an extended hydrophobic contact network that included Trp213 and resi ues from the surrounding site-I pocket (Figure 2). The proximity of the pose to Trp213 provides a structurally plausible explanation for the pronounced loss of intrinsic fluorescence. Conversely, the canonical site-II region in subdomain IIIA was not the targeted or supported pocket in this docking analysis.

**Figure 2.**
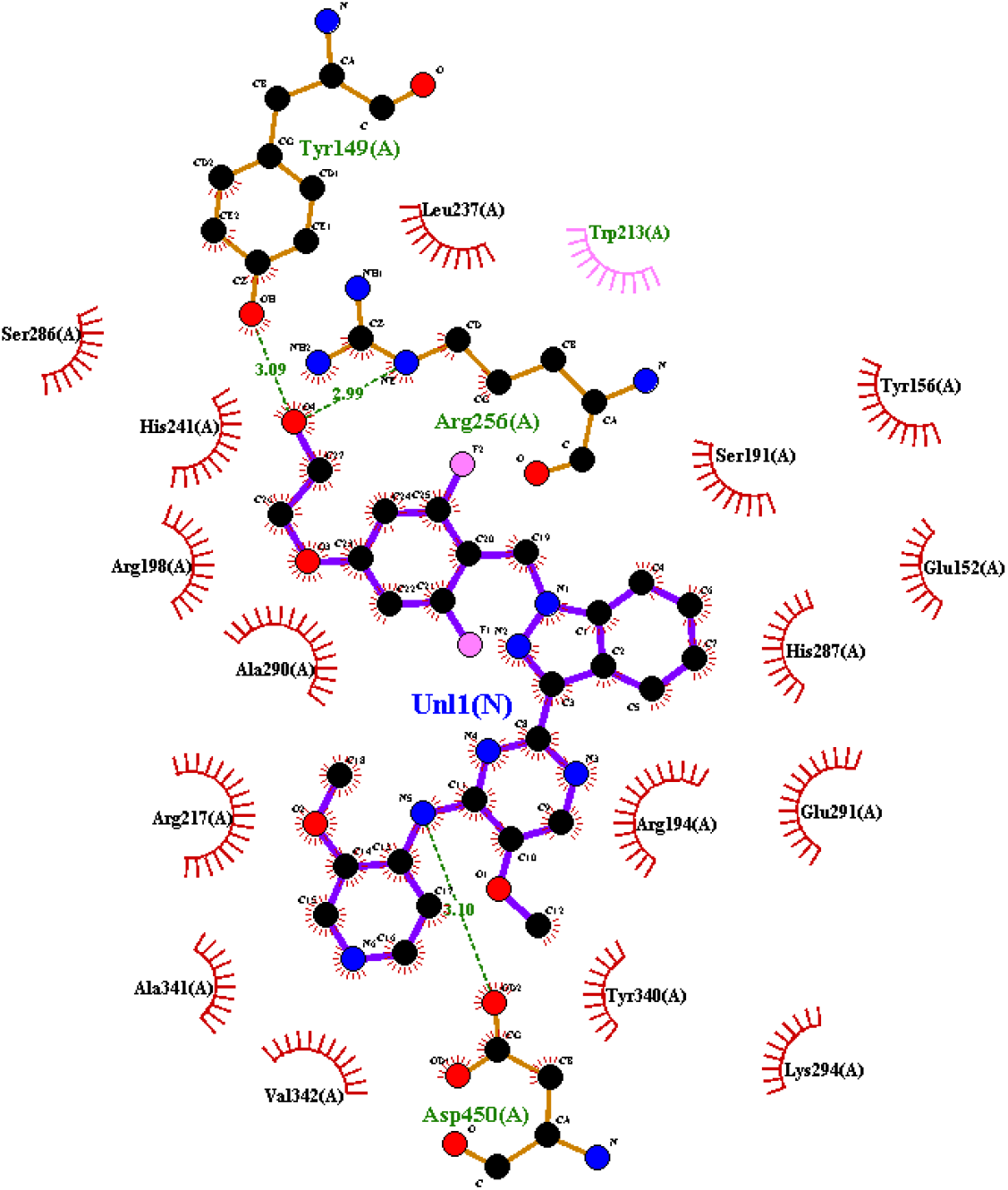
LigPlot+ interaction map for the top-ranked BAY-1816032 pose in BSA Sudlow site I. Green dashed lines denote predicted hydrogen bonds, with distances of 3.09 Å to Tyr149, 2.99 Å to Arg256, and 3.10 Å to Asp450. Red radial marks denote non-bonded contacts. The diagram uses the residue numbering and ligand label generated from the 4F5S docking file.

### 3.3 Added BSA reduces apparent BAY-1816032 potency in four mesothelioma models

Changing the basal serum concentration from 5% to 10% produced comparatively modest or cell-dependent changes in BAY-1816032 potency. In contrast, adding 2% (w/v) BSA to medium already containing 10% FBS produced a clear rightward shift in all four concentration–response curves (Figure 3; Table 1).

**Table 1.** BAY-1816032 IC□□ values under different serum and albumin conditions. Fold change = IC□□ in 10% FBS + 2% BSA divided by IC□□ in 10% FBS.

| Cell line | 5% FBS IC<sub>50</sub> (μM) | 10% FBS IC<sub>50</sub> (μM) | 10% FBS + 2% BSA IC<sub>50</sub> (μM) | Fold change vs 10% FBS |
| --- | --- | --- | --- | --- |
| H28 | 2.5 | 2.4 | 11.1 | 4.63 |
| H2052 | 2.0 | 3.6 | 22.0 | 6.11 |
| H2452 | 1.8 | 2.0 | 4.9 | 2.45 |
| AB12 | 2.9 | 2.4 | 11.1 | 4.63 |

**Figure 3.**
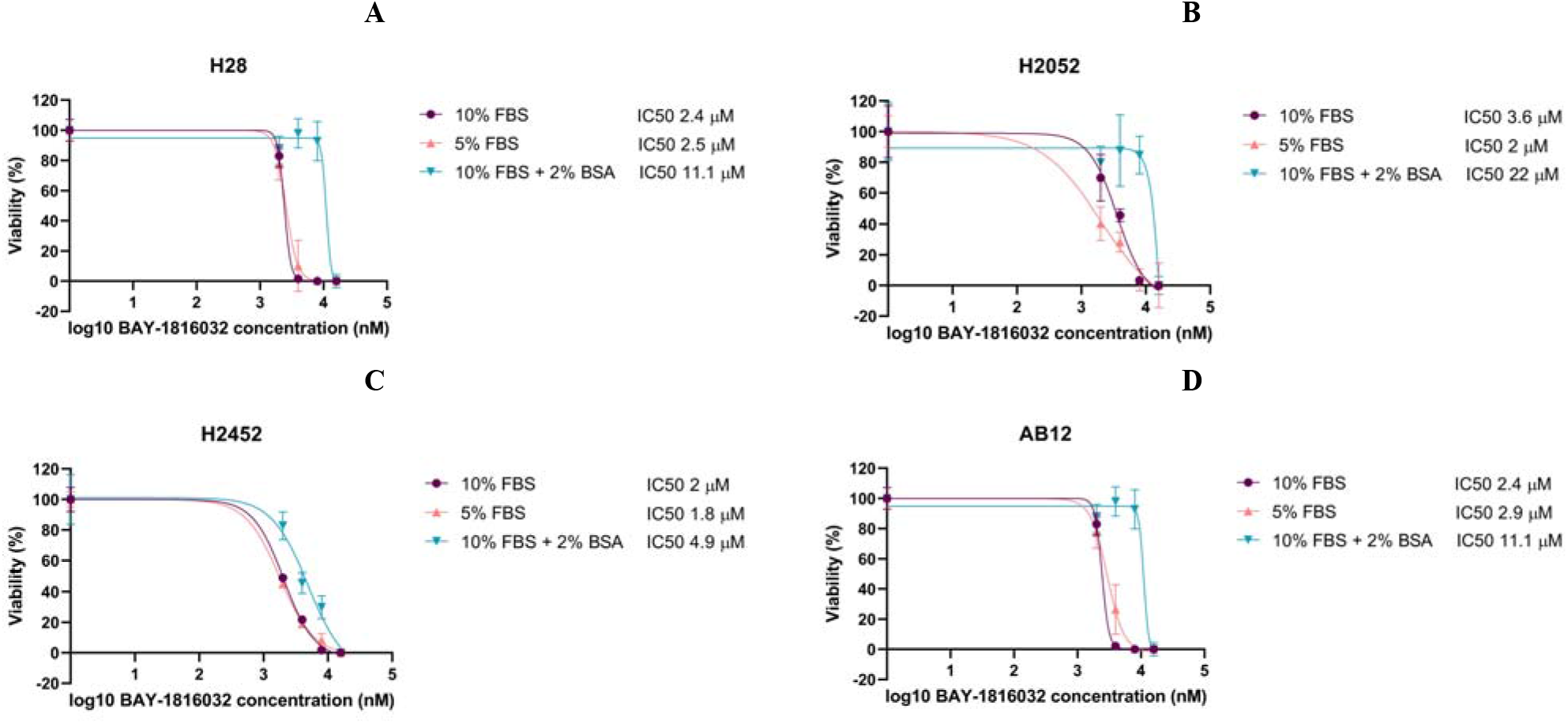
Effect of serum and added BSA on BAY-1816032 concentration–response curves in mesothelioma cells: (A) H28, (B) H2052, (C) H2452, and (D) AB12. Cells were evaluated in 5% FBS, 10% FBS, or 10% FBS supplemented with 2% (w/v; 20 mg/mL) BSA. The x-axis is logLL BAY-1816032 concentration in nM; the fitted IC□□ values are displayed in each panel.

The albumin-associated potency shift was largest in H2052 cells, in which the IC□□ increased from 3.6 to 22.0 µM (6.11-fold). H28 and AB12 each showed a 4.63-fold increase, from 2.4 to 11.1 µM, whereas H2452 showed a smaller but still substantial 2.45-fold increase, from 2.0 to 4.9 µM. The consistent direction across human and murine models supports a shared extracellular mechanism, while the range of fold changes indicates that cell-intrinsic factors may modulate the consequence of albumin association.

## 4. Discussion

The combined findings support a coherent model in which BAY-1816032 associates with albumin and the resulting protein reservoir lowers the concentration immediately available to enter mesothelioma cells. Three observations converge on this interpretation: pronounced quenching of intrinsic BSA fluorescence at low drug-to-protein ratios, a site-I docking pose close to Trp213, and a 2.45- to 6.11-fold loss of apparent cellular potency when 20 mg/mL BSA was added to serum-containing medium. The cellular experiment is particularly important because it connects the biophysical signal to a pharmacologically relevant consequence rather than treating albumin binding as an isolated spectroscopic phenomenon.

The docking result favors Sudlow site I, not site II. The pose occupies subdomain IIA, forms predicted hydrogen bonds with Tyr149, Arg256, and Asp450, and lies within the interaction environment of Trp213. BSA contains a single tryptophan residue, so perturbation of this region can strongly affect intrinsic fluorescence. Nevertheless, docking establishes a plausible structural hypothesis rather than occupancy in solution. A Vina score of −7.8 kcal/mol is not an experimental binding constant; converting it to an apparent micromolar value would import the scoring function’s approximations and should not be presented as KC or Kd. Competitive displacement by warfarin (site I marker) and ibuprofen (site II marker) would provide a direct test of the proposed site assignment [6].

The fluorescence response is compatible with complex formation, and the strong quenching at a 0.5:1 drug:BSA ratio is consistent with a substantial static component. However, fluorescence intensity alone cannot distinguish static quenching from dynamic quenching, inner-filter attenuation, or a combination of mechanisms. This distinction is especially important because BAY-1816032 absorbs in the near-UV region used to excite BSA [8]. Temperature-dependent Stern–Volmer measurements, fluorescence lifetime analysis, and absorbance-based inner-filter correction are therefore required before the mechanism is described as definitively static or a quantitative affinity is reported.

The pH experiment suggests that the BSA–BAY-1816032 optical response is enhanced under mildly acidic conditions. Albumin undergoes pH-dependent conformational adjustments, and pH can change both local residue states and ligand photophysics [7]. The present experiment does not resolve which contribution dominates. Its translational interpretation should therefore remain cautious: it demonstrates sensitivity over pH 7.1–7.7 but does not prove preferential drug release or retention in an acidic tumor microenvironment.

The MTT results offer an independent functional readout of albumin association. The consistent rightward shift after BSA addition in H28, H2052, H2452, and AB12 is most parsimoniously explained by a lower unbound fraction of BAY-1816032 in the extracellular medium. H2052 showed the largest shift and H2452 the smallest, despite the same albumin supplementation. Differences in drug uptake, efflux, target dependence, growth state, or intracellular handling could contribute to this variability. Without direct free-drug measurements or matched transporter/receptor experiments, the between-line variation should not be assigned to a specific albumin-uptake pathway.

This work also clarifies the observation that motivated the project. In the preceding serum-assay study, strong protein association necessitated 20% DMSO for quantitative extraction before HPLC–fluorescence analysis [8]. The present BSA and cell-based results identify albumin as a plausible major contributor to that matrix behavior. The validated analytical assay can now be repurposed for centrifugal-ultrafiltration experiments by measuring BAY-1816032 in the protein-retained and filtrate fractions, with appropriate nonspecific membrane-binding and mass-balance controls. Such measurements would convert the current mechanistic model into a quantitative unbound-fraction profile.

From a development perspective, albumin binding need not be uniformly unfavorable. Reversible association may increase circulating residence and solubility while reducing short-term free exposure. The practical issue is whether release is sufficiently rapid and whether delivery systems can preserve tumor exposure. The current findings therefore provide a rational baseline for later evaluation of liposomal or other carrier formulations under physiologically relevant albumin conditions, but formulation claims are beyond the evidence presented here.

## 5. Limitations

The study has four principal limitations. First, the fluorescence dataset lacks inner-filter correction, lifetime measurements, and temperature-dependent Stern–Volmer analysis; it therefore supports interaction but not a definitive quenching mechanism or experimental affinity constant. Second, docking was targeted to site I and was not complemented by an equivalent blinded or whole-protein comparison of sites I and II. Third, direct equilibrium binding and release were not measured by ultrafiltration, equilibrium dialysis, or an orthogonal biophysical method. Fourth, although the MTT study used six replicate wells per concentration with condition-specific blanks and controls, independent biological experiments and confidence-interval-based comparisons of the complete 4PL curves are required to establish biological reproducibility and formal statistical significance. These limitations do not negate the consistent albumin-dependent potency shift, but they define the experiments required for quantitative validation.

## 6. Conclusions

BAY-1816032 produces marked quenching of intrinsic BSA fluorescence, shows a predicted binding pose in Sudlow site I near Trp213, and loses apparent cellular potency when extracellular BSA is increased. Added BSA raised the BAY-1816032 IC□□ by 2.45- to 6.11-fold across four mesothelioma models, supporting the conclusion that albumin association reduces the immediately bioavailable drug fraction in vitro. The most defensible current interpretation is therefore qualitative and mechanistic: BAY-1816032 associates strongly enough with albumin to alter cellular exposure, while the binding constant, kinetic release behavior, and definitive site occupancy remain to be measured directly.

## Acknowledgments

This publication was created by benefiting from the 2236-B Marie Skłodowska-Curie Actions (MSCA) COFUND Scholarship Programmes Contribution Fund Programme (TÜBİTAK project ID: 123C460) and the Horizon Europe MSCA COFUND Postdoctoral Programme (EU project ID: 101126492). Views and opinions expressed are, however, those of the author(s) only and do not necessarily reflect those of the European Union or TÜBİTAK. Neither the European Union nor the granting authority can be held responsible for them. The authors gratefully acknowledge Bayer AG (Leverkusen, Germany) and the Structural Genomics Consortium (SGC, Frankfurt, Germany) for kindly providing BAY-1816032 used in this study. We especially express our deepest gratitude to Dr. Claudia Tredup for her careful coordination and support throughout the process of obtaining the compound. The authors also thank Ibrahim Oguzhan Tarman (IBG-FARMA, İzmir Biomedicine and Genome Center) for his valuable assistance in operating the RP-HPLC-FLD instrumentation.

## Funding

Funding information is included in the Acknowledgments section.

## Conflict of interest

The authors declare no financial competing interests.

## CRediT authorship contribution statement

Fazel Gorjipour: Conceptualization, methodology, investigation, formal analysis, data curation, visualization, writing – original draft, funding acquisition. Ece Çakiroglu: Conceptualization, methodology, investigation, formal analysis, data curation, visualization, and reviewing final proof. Şerif Şentürk: Conceptualization, supervision, resources, software, project administration, funding acquisition, writing – review and editing.

## Declaration of generative AI and AI-assisted technologies in the manuscript preparation process

During the preparation of this work, the authors used ChatGPT to improve text readability and structure. After using this tool, the authors reviewed and edited the content as needed and take full responsibility for the content of the article.

## Data availability

The data supporting the findings of this study are contained within the article and its figures. Additional raw data are available from the corresponding author upon reasonable request.

## Notes

### Competing Interest Statement

The authors have declared no competing interest.

